# Gene editing can generate fragile bivalents in mouse oocytes

**DOI:** 10.1101/350272

**Authors:** Marion Manil-Ségalen, Małgorzata Łuksza, Joanne Kannaan, Véronique Marthiens, Simon I.R Lane, Keith T. Jones, Marie-Emilie Terret, Renata Basto, Marie-Hélène Verlhac

## Abstract

Mouse female meiotic spindles assemble from acentriolar MTOCs (aMTOCs) that fragment into discrete foci. These are further sorted and clustered to form spindle poles, thus providing balanced forces for faithful chromosome segregation. To assess the impact of aMTOCs biogenesis on spindle assembly, we genetically induced their precocious fragmentation in mouse oocytes using conditional overexpression of Plk4, a master MTOC regulator. Excessive microtubule nucleation from these fragmented aMTOCs accelerated spindle assembly dynamics. Prematurely formed spindles promoted the breakage of three different fragilized bivalents, generated by the presence of recombined Lox P sites. Reducing the density of microtubules diminished the extent of chromosome breakage. Thus, improper spindle forces can lead to widely described yet unexplained chromosomal structural anomalies with disruptive consequences on the ability of the gamete to transmit an uncorrupted genome.

## Introduction

The microtubule (MT) spindle enables equal segregation of chromosomes between daughter cells during mitosis and meiosis. Checkpoints monitor the attachment of each chromosome to the spindle apparatus to ensure that this process is faithful (Etemad and Kops, 2016; Touati and Wassmann, 2016). Even though numerical errors induced by defects in spindle assembly have been extensively studied in mitosis and meiosis, not much is known on the mechanisms or the origins of structural errors arising during M-phase (Brunet et al., 2003; Kolano et al., 2012; Nagaoka et al., 2011; Sacristan and Kops, 2015). However, a link between abnormal mitosis and DNA damage has been established (Ganem and Pellman, 2012). For example, prolonged mitosis is often related to DNA damage (Rieder and Palazzo, 1992; Rieder and Maiato, 2004). Merotelic attachments, in which one kinetochore of a chromosome is attached to both spindle poles, are common in tumour cells with extra-centrosomes (Ganem et al., 2009). These attachment errors cause chromosome mis-segregation at mitotic exit (Cimini et al., 2001), and subsequent formation of micronuclei and chromothripsis in daughter cells (Cimini et al., 2001; Crasta et al., 2012; Zhang et al., 2015).

In the absence of canonical centrosomes, which are made from a pair of centrioles and pericentriolar material (PCM), meiotic spindles form with an important contribution from chromosome-mediated microtubule nucleation pathways. Thus MT assembly is enhanced in the vicinity of meiotic chromosomes (Heald et al., 1996; Dumont et al., 2007; Dumont and Desai, 2012; Bennabi et al., 2016). In mouse oocytes, PCM aggregates remain despite centriole loss, a feature that describes them as acentriolar MicroTubule Organizing Centers (aMTOCs). It is unclear how aMTOCs assemble in the absence of centrioles but they largely contribute to meiotic spindle assembly (Maro et al., 1985; Dumont et al., 2007; Schuh and Ellenberg, 2007). In fully grown prophase I-arrested oocytes, aMTOCs constitute typically 1-3 large foci (Luksza et al., 2013). At meiotic entry, these foci stretch around the nuclear envelope and fragment in a dynein-and MT-dependent manner (Luksza et al., 2013; Clift and Schuh, 2015). The importance of such timely reorganization of aMTOCs has not been addressed.

Plk4 is the master regulator of centriole duplication (Bettencourt-Dias et al., 2005; Habedanck et al., 2005), and is also essential for meiotic and mitotic acentriolar spindle assembly in oocytes and early mouse embryos by stimulating MT growth (Bury et al., 2017; Coelho et al., 2013). In the present study, we took advantage of the role of Plk4 to perturb aMTOCs organization in oocytes (Luksza et al., 2013). Transgenic overexpression of Plk4 (Marthiens et al., 2013) in oocytes during their growth phase in the ovary led to precocious aMTOC fragmentation before nuclear envelope breakdown (NEBD). It also led to earlier spindle bipolarisation associated with greater MT nucleation. Following NEBD, oocytes completed meiosis I. However one bivalent broke into two pieces, consistent with breaks occurring near the recombined Lox P sites present in the genome of three different mouse lines. Crucially, a decrease in MT density diminished the efficiency of bivalent breakage. Altogether, our work suggests that increasing MT density and probably forces exerted on fragilized chromosomes can lead to bivalent breakage during meiosis I, as shown here for three different chromosomes. This in turn leads to mis-segregation of chromosome fragments. Chromosome rearrangements and fragment losses are relatively frequent in human embryos and lead to severe pathologies. This is the case for intellectual disability syndromes such as the Wolf-Hirschhorn due to a 4p deletion (Wieczorek et al., 2000), the 9p deletion (Okten et al., 2009) or the 6q terminal deletion (Stria et al., 2006). We hypothesize that our work may identify one mechanism by which structural chromosome anomalies can arise during meiosis.

## Results

### Plk4 overexpression in growing oocytes causes premature aMTOC fragmentation

As previously published (Bury et al., 2017), we confirmed that Plk4 is present in mouse oocytes. For this we labelled endogenous Plk4 by immunofluorescence in control oocytes blocked in prophase I of meiosis (Fig. 1A bottom left panel). By co-staining with the PCM component, Pericentrin (Fig. 1A top left panel), we showed that Plk4 localizes to aMTOC arguing that it is not a limiting factor for centriole assembly in oocytes.

**Fig. 1.**
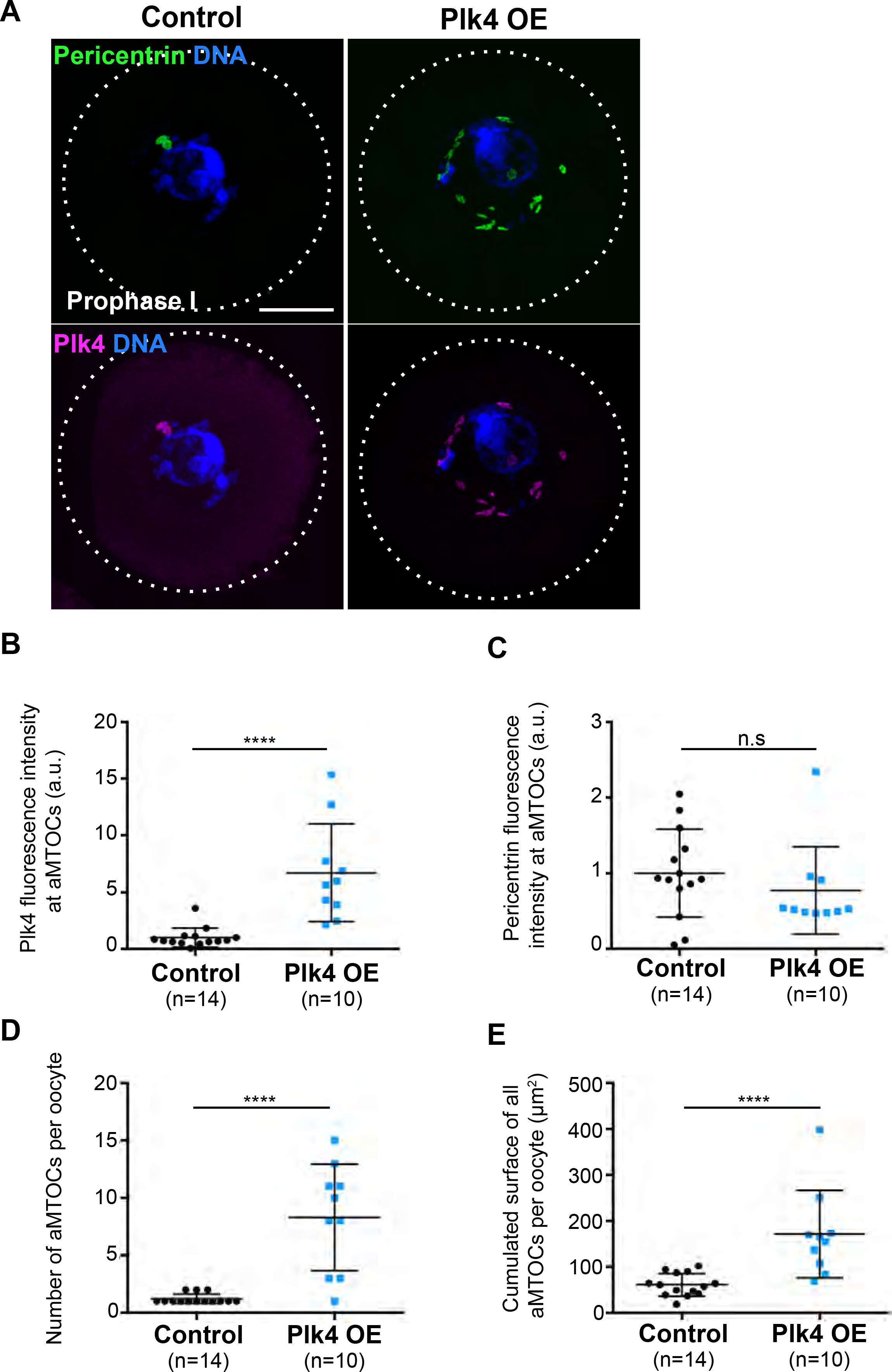
Plk4 OE induces aMTOC precocious fragmentation. **(A)** Immunofluorescent co-staining of Plk4 (pink) and Pericentrin (green) in control (left panels) and Plk4 OE (right panels) oocytes observed in prophase I. Scale bar is 10 μm. **(B)** Measure of Plk4 fluorescence intensity exclusively at nucleus associated aMTOCs from control (black dots) and Plk4 OE oocytes (blue squares) as observed in (A), arbitrary units, p<0.0001. **(C)** Measure of Pericentrin fluorescence intensity exclusively at nucleus associated aMTOCs from control (black dots) and Plk4 OE (blue squares) oocytes as observed in (A), arbitrary units, p=0.2125. **(D-E)** Quantitative analysis of aMTOCs number per oocyte **(D)** and their cumulated surface per oocyte (E) in control (black dots) and Plk4 OE (blue squares) oocytes in prophase I as observed in (A). p<0.0001 for number and surfaces. In (**B-E**) statistical tests used are two-tailed Mann-Whitney and n corresponds to the number of oocytes.

Conditional Plk4 overexpression in oocytes was achieved by crossing a line expressing the Cre-recombinase under the ZP3 promoter with a line containing a random insertion of a chicken β-actin promoter (CAG) fused to a floxed stop-codon and to mCherry-Plk4 (Fig. S4A, referred as Plk4 OE; Marthiens et al., 2013). The ZP3 promoter is active after birth, after homologous chromosomes have recombined and formed chiasmata and only during oocyte growth (Lewandoski et al., 1997). In this way, we obtained excision of the stop-codon and expression of the m-Cherry-Plk4 transgenic construct exclusively in oocytes during their growth period at puberty.

In both Plk4 OE prophase I arrested oocytes (from Plk4^flox/wt^; Cre^+^ female mice), and control oocytes (coming either from Plk4^flox/wt^; Cre^-^ or Plk4^wt/wt^; Cre^+^ female mice), Plk4 was detected on aMTOCs (Fig. 1A bottom panels). However, Plk4 OE oocytes displayed modified aMTOCs characteristics with numerous small foci in the vicinity of the nucleus (Fig. 1A right panels). In Plk4 OE oocytes, the amount of Plk4 at aMTOCs was more than 6x higher than in controls (Fig. 1B) while the amount of Pericentrin at aMTOCs remained unchanged (Fig. 1C). Due to the relative size of the oocyte and the aMTOCs, the local aMTOCs signal corresponds to only 0.1% of the signal in the entire oocyte. It is thus not so surprising that no detectable increase in the total amount of either Pericentrin or Plk4 was observed in whole Plk4 OE oocytes compared to controls (Fig. S1A and S1B). These data argue that exogenous Plk4 preferentially accumulates on aMTOCs and is present there in large excess.

To describe the changes in the organization of aMTOCs we performed 3D quantitative analysis of competent, fully grown Plk4 OE oocytes in prophase I (Fig. 1D-E). The number and the cumulated surfaces of all aMTOCs were both significantly higher in Plk4 OE compared to controls. These observations suggested that aMTOCs were more fragmented in Plk4 OE oocytes.

### aMTOC fragmentation occurs at the end of the prophase I block

We used 3D structured illumination microscopy (3D-SIM) to look more closely at the detailed morphology of PCM in prophase I arrested Plk4 OE oocytes, as it has been done in centriolar cells (Fu and Glover, 2012; Lawo et al., 2012; Mennella et al., 2012; Sonnen et al., 2012). In controls, oocytes display one major aMTOC, harbouring a globular shape with the Pericentrin matrix displaying multiple cavities filled with Plk4 (Luksza et al., 2013). In contrast, fully-grown Plk4 OE oocytes, competent to resume meiosis I, contained multiple aMTOCs that assembled into thread-like structures when running parallel to the nuclear envelope (Fig. S2A, right panels). The Pericentrin internal cavities were no longer detected in these long thread-like structures, indicating a dramatic change in PCM organization.

The super-resolution microscopy data, showing an absence of increase in Pericentrin labelling together with the 3D quantifications of aMTOCs organisation argue for a premature fragmentation of the PCM in Plk4 OE oocytes (Fig. S2B). This fragmentation occurs at the end of oocyte growth, in prophase I, as incompetent Plk4 OE oocytes present only one globular Pericentrin matrix, filled with Plk4, as in controls (Fig. S2A, left panels and S2B; Luksza et al., 2013). Thus, even if induced early on during the growth phase, Plk4 overexpression results in major changes in the organization of aMTOCs only at the end of the prophase I block, once the oocyte reaches competency to resume meiosis I.

### Overexpression of Plk4 during the growth phase of the oocyte is needed for aMTOCs fragmentation

The fragmentation of aMTOCs occurred at the end of the oocyte growth phase, thus we tested whether a similar phenotype could be reproduced by directly injecting the cRNA encoding for mCherry-Plk4 into fully-grown oocytes in prophase I (Fig. 2A). When the mCherry-Plk4 cRNA is injected, much higher levels of Plk4 are reached than with Plk4 OE, both globally and associated to aMTOCs (Fig. 2B, C and D). In the case of the cRNA, the total number of aMTOCs per oocyte was higher than in controls but was comparable to the number observed in the transgenic line overexpressing Plk4 (Fig. 2B and E). However, instead of several aMTOCs presenting width ranging from 1 to 5 μm^2^ as in Plk4 OE oocytes (Fig. 2F), in each injected oocyte we detected one large aMTOC and multiple tiny foci of mCherry-Plk4 (smaller than 1 μm^2^; compare left versus right panel on Fig. 2B and see size quantification on Fig. 2F). Although we do not know how these very small aMTOCs are generated, the large aMTOC is the dominant site for MT nucleation in these oocytes (Fig. 2G left panel). This is strikingly different from what can be observed in the Plk4 OE transgenic prophase I oocytes where all fragments of aMTOCs show significant MT nucleation (Fig. 2G right panel). Altogether our results suggest that Plk4 overexpression during the growth phase of oocytes (via transgenic Plk4 OE), but not when limited to the competent/fully-grown stage (via cRNA injection), leads to premature fragmentation of aMTOCs into smaller functional units able to promote significant MT nucleation.

**Fig. 2.**
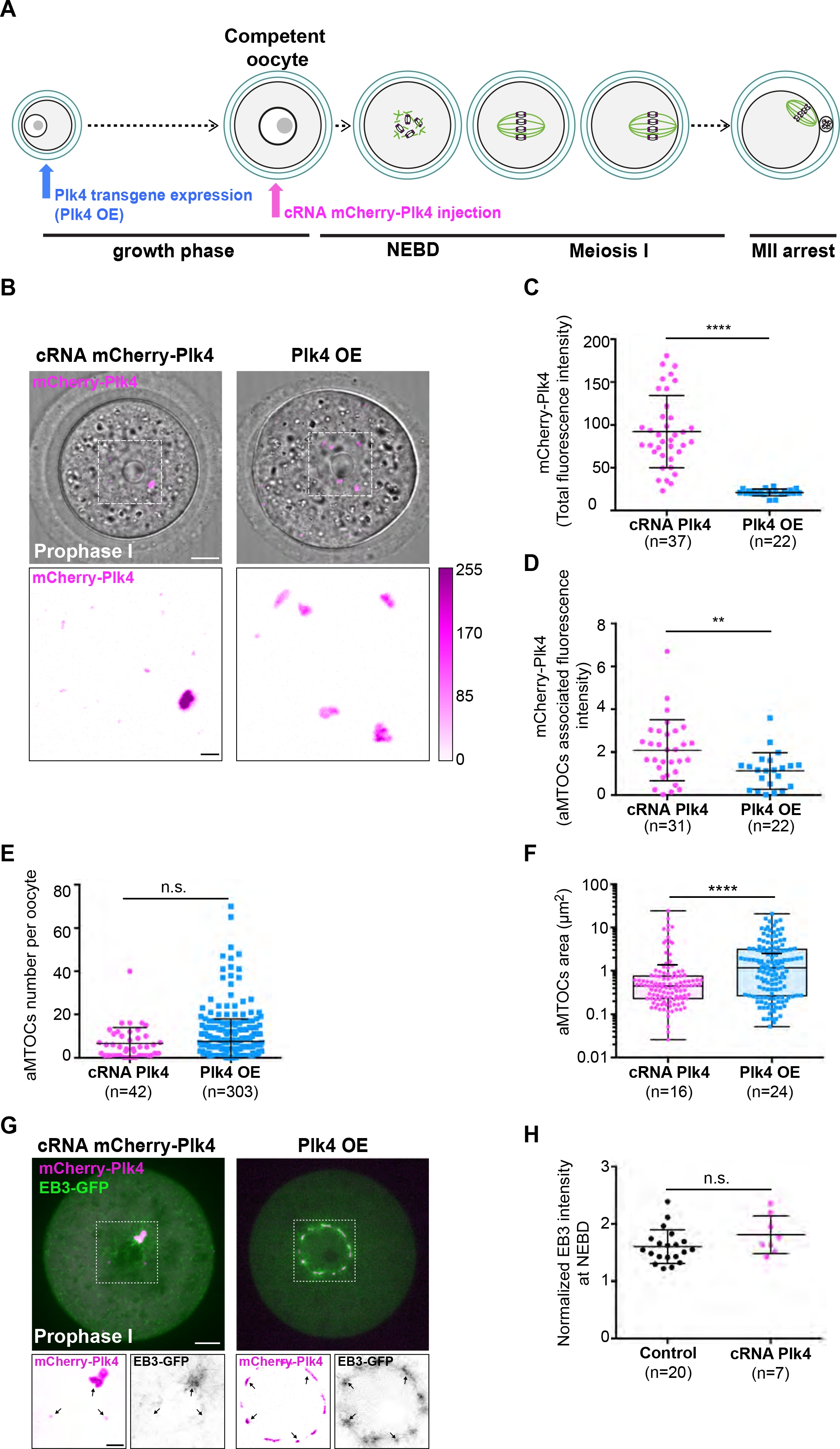
Over-expression of Plk4 during oocyte growth induces aMTOC fragmentation. **(A)** Schematic of the two methods for achieving mCherry-Plk4 OE in mouse oocytes: either via a transgenic approach allowing Plk4 OE at the beginning of the growth phase (in blue) or by mCherry-Plk4 cRNA injection allowing Plk4 OE after the growth phase (in pink). mCherry-Plk4 cRNA was injected into competent/fully-grown control oocytes and expressed 3 h before live imaging. Plk4 OE oocytes were imaged directly upon isolation. Chromosome (gray); kinetochores (pink); MTs (green). NEBD, Nuclear Envelope BreakDown; MII, Metaphase II. **(B)** Upper panels correspond to the merge of transmitted light images with the mCherry-Plk4 fluorescent image in cRNA injected (left panel) versus transgenic Plk4 OE (right panel) oocytes observed in prophase I (mCherry-Plk4, magenta). The white dotted square highlights the nuclear area and the mCherry-Plk4 signal from this region is displayed on the lower panels. The signal intensity lookup table is depicted on the right side. Scale bar is 10 μm (upper panels) or 2 μm (lower panels). **(C)** Total levels of mCherry-Plk4 expression measured as absolute signal intensity from oocytes in prophase I either cRNA injected (purple dots) or from Plk4 OE (blue squares) as observed in (B). p<0.0001 (two tailed t-test with Welch correction). (**D**) aMTOC-associated levels of mCherry-Plk4 overexpression measured as absolute signal intensity from oocytes in prophase I either cRNA injected (purple dots) or from Plk4 OE (blue squares). p=0.0046. **(E)** Quantitative analysis of aMTOCs number in oocytes injected with mCherry-Plk4 cRNA (purple dots) and from Plk4 OE (blue squares). p=0.8752. **(F)** aMTOCs area in oocytes injected with mCherry-Plk4 cRNA (purple dots) and from Plk4 OE (blue squares). Box and whisker plots represent the min and the max, boxes limits correspond to the 25^th^ and the 75^th^ percentiles. The long horizontal line is the median; the short horizontal line is the mean (p<0.0001). (**G**) EB3-GFP (green) is presented to visualise MT organisation in cRNA injected (left panels) versus Plk4 OE (right panels) prophase I oocytes. The EB3-GFP signal (green) is merged with the mCherry-Plk4 signal (magenta). The white dotted boxes highlight the nuclear regions. Insets are higher magnifications of signals from nuclear regions. In insets, the EB3-GFP signal appears black while mCherry-Plk4 appears magenta. The black arrows point towards aMTOCs. Scale bars are 10 μm (upper panels) and 5 μm (lower insets). **(H)** Normalized signal intensity of the EB3-GFP signal around chromosomes in oocytes observed at NEBD from controls (black dots) or injected with mCherry-Plk4 cRNA (pink dots). p=0.1358. In **(D-F)** and in **(H)**, the statistical tests used were two-tailed Mann-Whitney. In all graphs n is the number of oocytes.

### Fragmented aMTOCs have increased MT-nucleating capacity

To test the impact of precocious aMTOCs fragmentation, oocytes were then allowed to resume meiosis and local MT density was observed around the condensing chromosomes at the time of NEBD as done previously (Breuer et al., 2010). The intensity of EB3-GFP fluorescence in the vicinity of chromosomes was significantly higher (1.4x) in Plk4 OE oocytes indicating that they contained more MT than controls (Fig. 3A and B). Importantly, the increase in EB3-GFP intensity at NEBD correlated with the number of aMTOCs observed in prophase I (Fig. 3C). However such an increase in MT density at NEBD was not observed after overexpression of mCherry-Plk4 via cRNA injection into fully-grown oocytes (Fig. 2H), despite higher levels of overexpression reached by this approach (Fig. 2C and D). This result suggests that the extent of aMTOCs fragmentation in prophase I impacts on the MT density at meiosis resumption.

**Fig. 3.**
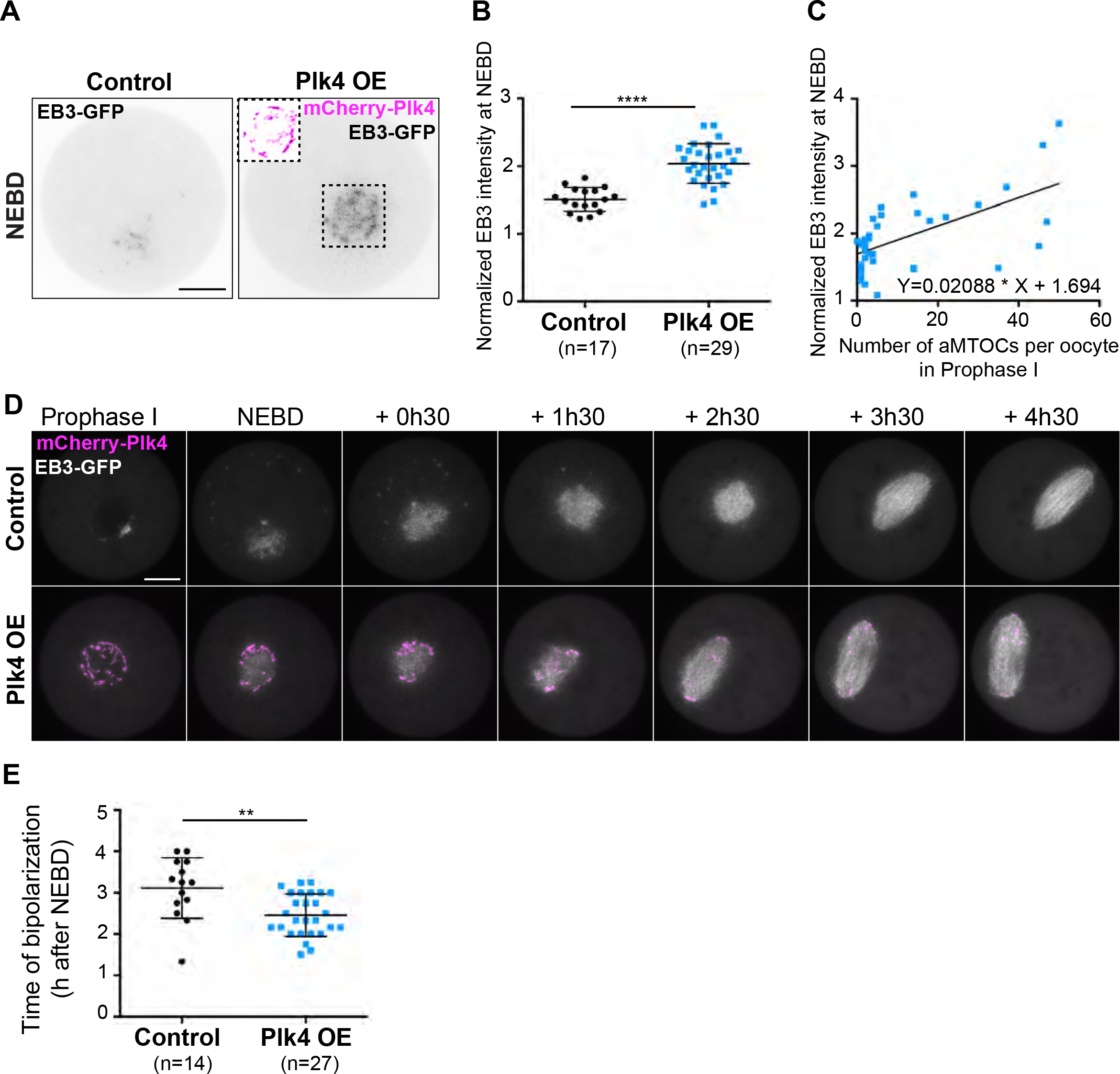
Fragmented aMTOCs have increased MT-nucleating activity at NEBD. **(A)** EB3-GFP expression in control (left panel) and Plk4 OE (right panel) oocytes observed at NEBD. The black dotted square shows the mCherry-Plk4 labelling (pink) that surrounds early spindle MT, visible only in Plk4 OE oocytes. EB3-GFP appears in gray levels. Scale bar is 10 μm. (B) Normalized signal intensity of EB3-GFP around chromosomes in control (black dots) and Plk4 OE (blue squares) oocytes observed at NEBD. The signal was measured in a Ø20 μm region around the chromosomes; total signal intensity was measured in a Ø70 μm region covering the whole oocyte. Normalized signal intensity is the ratio of local/total intensity measured for individual oocytes. p<0.0001. **(C)** Correlation between the number of aMTOCs in prophase I and normalized fluorescence intensity of EB3-GFP at NEBD (n=37). Normalized EB3-GFP fluorescence intensity measured at NEBD is plotted against the number of aMTOCs per oocyte in prophase I. **(D)** Early steps of spindle formation in control versus Plk4 OE oocytes expressing EB3-GFP. Oocytes were followed from prophase I exit until spindle bipolarization. EB3-GFP appears white and mCherry-Plk4 pink. Scale bar is 15 μm. Timing (expressed in hours and minutes) is relative to NEBD. **(E)** Timing of spindle bipolarization observed in living oocytes expressing EB3-GFP as presented in (D), p=0.0076. For (**B**) and (**E**), the statistical tests used are two tailed t-test with Welch correction. For (**B**) and (**E**), n corresponds to the number of oocytes.

To estimate the consequences of an increased MT density at NEBD, we followed the formation of meiotic spindles in live EB3-GFP injected oocytes (Fig. 3D and Supplementary Movie S1). In control oocytes, the entire meiosis I lasts around 9 h from NEBD until polar body extrusion (PBE). At early stages of meiosis resumption after NEBD, the newly assembled mass of MT self-organizes into a bipolar array around 3-4 h after NEBD (Dumont et al., 2007; Schuh and Ellenberg, 2007). As expected from previous work (Dumont et al., 2007; Kolano et al., 2012), increased density of MT observed in Plk4 OE oocytes resulted in earlier spindle elongation and bipolar axis formation (on average 45 min earlier in Plk4 OE oocytes compared to controls; Fig. 3D and E). This result was confirmed on fixed samples observed 4 h after NEBD and analysed by immunofluorescence (Fig. S3A and B). More oocytes (61.4% versus 26.1%) contained a robust bipolar spindle in Plk4 OE oocytes compared to controls at 4 h after NEBD (Fig. S3B). Thus, in Plk4 OE oocytes meiotic spindle bipolarity is established earlier than in controls.

### Plk4 OE oocytes are endowed with chromosome breakage

We then analysed the consequences of premature increased density of MT early on during meiosis I. Unexpectedly, Plk4 OE oocytes frequently contained 21 individual chromosome fragments, instead of the normal 20 bivalent complement of mouse. Remarkably, we observed the appearance in live cells of a round chromosome, smaller than all others that oscillated back and forth across the metaphase plate, suggesting it was not correctly bi-oriented (Fig. 4A, black arrows and Supplementary Movie S2). Indeed, this small DNA fragment had only one pair of kinetochores positive for Mad2-YFP (Fig. 4A right inset). We wanted to characterise better this breakage event, which has never been described previously in the first meiotic division without prior treatment with a DNA damaging agent (Collins et al., 2015; Lane et al., 2017; Marangos et al., 2015). Therefore, oocytes were fixed at NEBD +6h00, centromeres were labelled with CREST and 3D reconstruction used to count the number of bivalents in oocytes derived from two independent insertions of the same Plk4 transgene (transgenic line A and line B, see Methods, Fig. S4C and Supplementary Movie S3, S4 and S5). In the transgenic line A, we observed the breakage of one bivalent as depicted in Fig. 4A: one large and one small fragment, both of which harbouring a pair of centromeres labelled with CREST (Fig. 4B upper panel and Supplementary Movie S4, compare with Movie S3). The pattern of breakage was very reproducible within this line (Fig. 4C showing various examples artificially highlighted in blue). In line B, we also observed the breakage of a single bivalent, but different from line A, generating a fragment of DNA lacking CREST signal (Fig. 4D and 4E and Supplementary Movie S5) and much smaller than any fragment detected in oocytes from line A (Fig. 4F). Without any centromere and so devoid of the ability to assemble a kinetochore, this smaller fragment was not retained in the meiotic spindle (Fig. 4D left picture and other examples of the small fragment artificially highlighted in green in Fig. 4E). The frequency at which we could observe chromosome breakage in the two lines differed: it was more frequent in line A (27.0%) than in line B (10.2%; Fig. 4G and summarized in Fig. S4C). Differences in the rate of chromosome breakage between the two strains were not a result of different overexpression levels of Plk4 as no difference in mCherry-Plk4 expression was found at aMTOCs between the two lines (Fig. S5A and S5B).

**Fig. 4.**
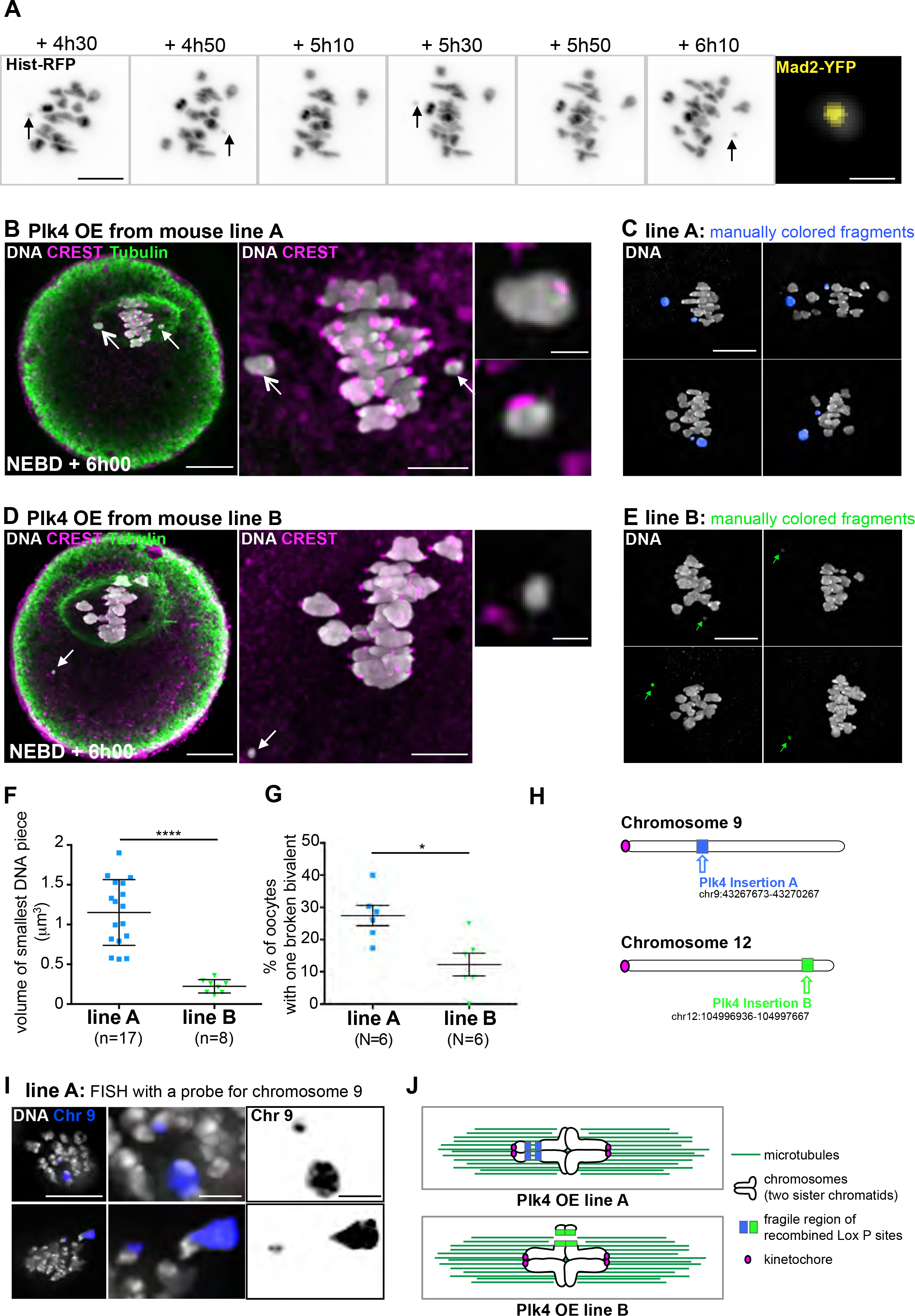
Evidence for chromosome breakage in two transgenic lines over-expressing Plk4. **(A**) Time-lapse images of chromosomes labelled with histone-RFP followed in Plk4 OE (line A) during meiosis I. Black arrows point at the smallest chromosome fragment moving back and forth the metaphase plate. Chromosomes appear in gray levels. Timing (expressed in hours and minutes) is relative to NEBD. Scale bar is 10 μm. The right panel shows the smallest DNA fragment (gray) at a higher magnification harbouring a Mad2-YFP signal (yellow) at NEBD + 6h10. Scale bar is 1 μm. **(B, D)** Immunofluorescent co-staining of DNA (white), CREST (pink) and Tubulin (green) in Plk4 OE line A (**B**) and line B (**D**) oocytes observed at NEBD + 6h00. White arrows point at the chromosome pieces. Higher magnification of DNA pieces is shown on the right panels: the upper right panel corresponds to the largest one and the lower right panel to the smallest one. Scale bar is 20 μm in the left panels, 5 μm in the middle panels and 1 μm in the right panels. **(C, E)** Examples of fixed oocytes with broken bivalents in line A (**C**) and line B (**E**) observed at NEBD + 6h00. Broken bivalents have been artificially coloured in blue for line A (**C**) and in green for line B (**E**). Scale bar is 5 μm. **(F)** Quantitative analysis of the volume of the smallest chromosome fragment in oocytes from Plk4 OE lines A and B observed at NEBD + 6h00. p<0.0001 (two tailed t-test with Welch correction). n is the number of oocytes. **(G)** Quantification of the percentage of oocytes with one broken bivalent in Plk4 OE line A and B observed at NEBD + 6h00. p= 0.0104 (Fisher exact test). N is the number of mice. **(H)** Scheme representing the positions of the two mCherry-Plk4 transgenes: the insertion (in blue) is closer to the centromeric region (in pink) of chromosome 9 for line A and the insertion (in green) is closer to the telomeric region of chromosome 12 for line B. **(I)** Two examples of FISH labelling of chromosome 9 (blue) in line A observed at NEBD + 6h00. 36 oocytes were collected from 3 mice. Among them, 16.67% had a broken bivalent. These broken bivalents were always labelled by the FISH probe for chromosome 9. DNA is labelled in gray. The FISH signal appears in blue. Scale bar is 10 μm for the left upper panel. Higher magnifications are shown in the middle and right upper panels. Scale bar is 2 μm. Chromosome fragments labelled by the FISH probe appear in black on the right panels. (**J**) Proposed model of bivalent breakage occurring in the two Plk4 OE lines. MTs appear in dark green. Kinetochores are in pink. The mCherry-Plk4 transgene insertion sites appear in blue (line A) and green (line B).

Since the breakage occurred only on one bivalent, in a specific manner for both transgenic lines, we hypothesized that it took place at the insertion site of the mCherry-Plk4 transgene-this being the only major difference between both lines. The site of insertion was sequenced for both lines. In line A, it was found to be close to the centromeric region of chromosome 9, theoretically generating a small 45 Mb DNA fragment if the cut site occurs in the region of the transgene. In line B it was close to the telomeric region of chromosome 12, which would generate a very small 15 Mb acentric DNA fragment if the cut occurs in the region of the transgene (Fig. 4H and S4C). These sites of insertion are fully consistent with the fragments observed in both lines: two fragments containing centromeres for line A (Fig. 4B), and a smaller acentric fragment for line B (Fig. 4D).

To support the sequencing data, we performed fluorescent in situ hybridization (FISH) experiments on oocytes from line A. The chromosome fragments corresponded to the genome location of transgene insertion, as 100% of the observed broken chromosomes were labelled with a FISH probe for chromosome 9 (Fig. 4I). Chromosome breaks were never observed in oocytes coming from Plk4^flox/wt^; Cre^-^ mice or Plk4^wt/wt^; Cre^+^ female mice (our negative controls that do not overexpress Plk4) nor in the mCherry-Plk4 cRNA injected oocytes (Fig. S3C). Furthermore, we know that recombination and repair of the mCherry-Plk4 transgene did happen in all Plk4 OE oocytes from both lines, since 100% of them were positive for mCherry-Plk4. Therefore it can be excluded that the break took place early on during oocyte growth. Our data strongly support the view that the chromosome breakage occurs at or near the site of the recombined transgenic insertion at late stages of oocyte growth or even later during the division (Fig. 4J).

### Chromosome breakage can occur at any Cre-recombined Lox P site provided aMTOCs are fragmented

We then tested whether the creation of another fragile site, different from the transgenic Plk4 insertion, could also be associated with breakage of one bivalent. For this we crossed Plk4 line A^flox/wt^; Cre^+^ mice with MyoX^flox/wt^; Cre^-^ mice. The MyoX^flox/wt^; Cre^-^ line has been constructed by insertion of two Lox P sites between exon 26 and exon 30 of the Myosin X endogenous locus in chromosome 15 (Fig. S4B). These mice are fertile and display no obvious phenotype in their oocytes (data not shown and Fig. 5 lower panels). Oocytes coming from the Plk4 line A^flox/wt^; Cre^+^; MyoX^flox/wt^ females were fixed at NEBD + 6h00 (as done in Fig. 4B and 4D) and their chromosomes were counted. 13.6% of these oocytes displayed a broken bivalent resembling the one observed in Plk4 line A (Fig. 5A, upper panels, blue arrows), while 15.6% presented a new type of break: a large piece of DNA, with no centromere (Fig. 5A, lower panels, orange arrow and Supplementary Movie S6). We did not observe oocytes presenting simultaneously the two types of break. Their frequency might be too low and we did not look at enough oocytes coming from this double cross (Fig. 5B). The volume of the new acentric fragment was significantly larger than the smallest DNA piece observed in both line A and line B (Fig. 5C and D). We interpret this new type of break as a breakage at the MyoX locus on chromosome 15 (Fig. 5E and 5F), since it is consistent with the larger size of the acentric fragment as well as with the potential chiasma location on this chromosome (Morgan et al., 2017). Acentric fragments of chromosomes are not retained in the spindle (see Fig. 4D). Remarkably, we observed 2/3 of oocytes with the large acentric fragment close to the metaphase plate (Fig. 5D left picture), while 1/3 was away from the spindle (Fig. 5D right picture). This observation could indicate that the fragile bivalent is cut into two pieces while meiosis I progresses. Importantly, no chromosome break was observed in control oocytes from Plk4 line A^flox/wt^; Cre^-^; MyoX^flox/wt^ or Plk4 line A^wt/wt^; Cre^+^; MyoX^flox/wt^ females (Fig. 5A lower panels). This indicates that the break occurs only when two conditions are met: Plk4 overexpression associated with aMTOCs fragmentation and recombined Lox P sites. These data argue that sites breaking under tension can be created in the genome, outside the Plk4 insertion.

**Fig. 5.**
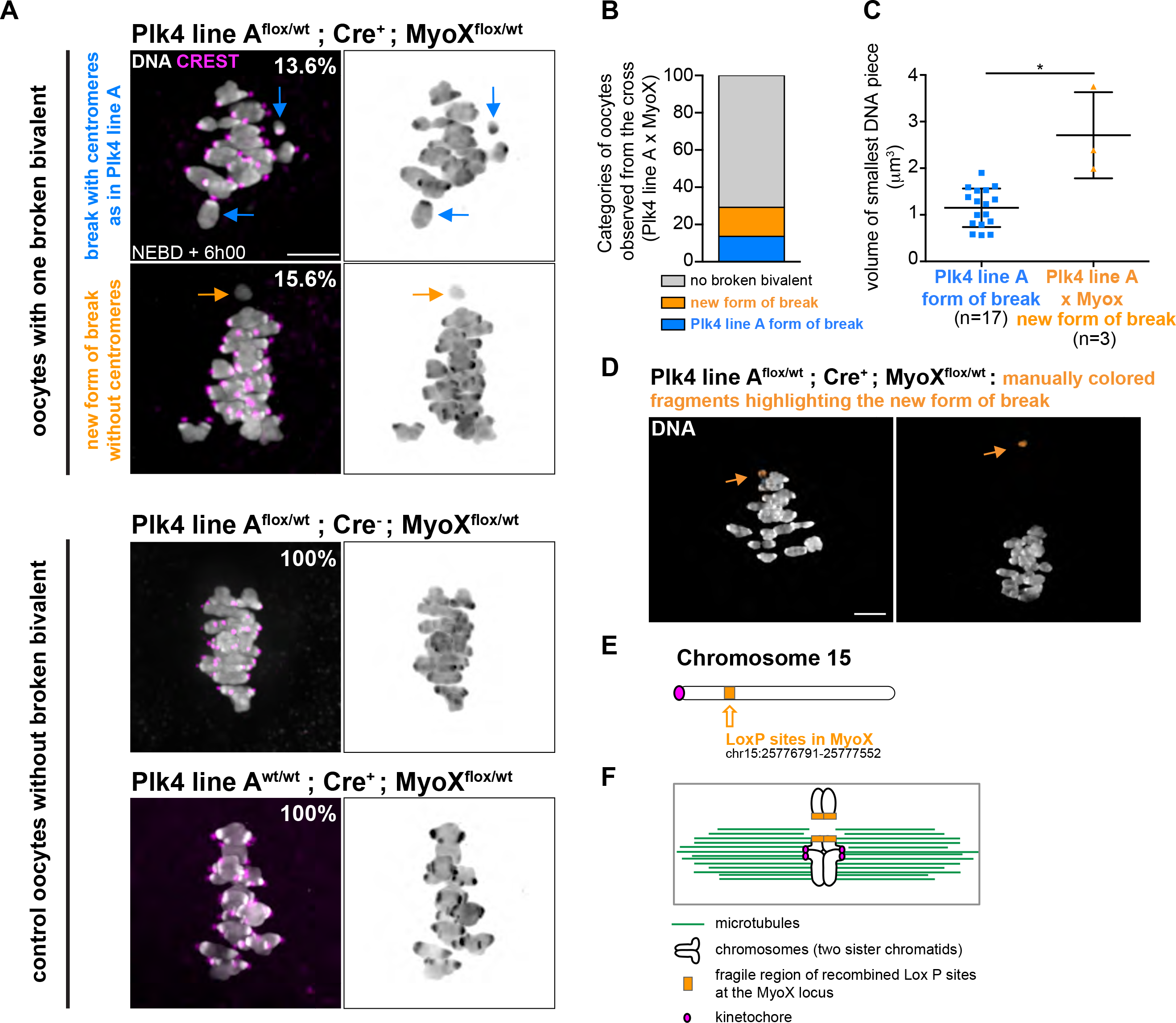
The breakage is not specific of the Plk4 transgenic insertion. **(A)** Oocytes coming from a Plk4 line A^flox/wt^; Cre^+^; MyoX^flox/wt^ cross display both Plk4 line A type of break (13.6% of oocytes, upper panels, see blue arrows) and a new form of break producing a fragment with no centromeres (15.6 % of oocytes, middle panels, see orange arrows). DNA is in gray levels, CREST in magenta. The two lower panels are showing control oocytes (Plk4 line A^flox/wt^; Cre^-^; MyoX^flox/wt^ and Plk4 line A^wt/wt^; Cre^+^; MyoX^flox/wt^). The percentage of oocytes without broken bivalent from these controls is indicated on the pictures of the two left lower panels. Scale bar is 5 μm. **(B)** Percentages of oocytes with the different types of break in the Plk4 line A^flox/wt^; Cre^+^; MyoX^flox/wt^ cross (on n=32 oocytes). **(C)** Quantitative analysis of the volume of the smallest chromosome fragment in oocytes from Plk4 OE lines A (blue squares) and oocytes from the Plk4 line A^flox/wt^; Cre^+^; MyoX^flox/wt^ cross (orange triangles) observed at NEBD + 6h00. p=0.0121 (Mann-Whitney test). n is the number of oocytes. **(D)** Examples of oocytes with the new type of broken bivalent in Plk4 line A^flox/wt^; Cre^+^; MyoX^flox/wt^ cross observed at NEBD + 6h00. Broken bivalents have been artificially coloured in orange. Scale bar is 5 μm. **(E)** Scheme representing the position of the MyoX LoxP sites (orange) on chromosome 15. **(F)** Proposed model of the new bivalent breakage occurring in Plk4 line A^flox/wt^; Cre^+^; MyoX^flox/wt^ cross. MTs are in green. Kinetochores are in pink. The LoxP sites insertion region appears in orange.

### Extent of bivalent breakage correlates with the extent of aMTOCs spreading and MT density

Plk4 OE oocytes have fragmented aMTOCS, nucleate more microtubules and thus present an accelerated spindle formation. All this could cause changes in forces applied on chromosomes during meiosis I and favour chromosome breakage. We went on to characterize further the relationship between spindle dynamics and the generation of broken bivalents.

First, we compared line A and line B. As mentioned previously, more chromosome fragments are observed in line A than in line B (Fig. S4C). Consistently, we observed that the extent of aMTOCs fragmentation in prophase I was higher in line A than in line B despite them having the same level of overexpression at the end of oocyte growth (Fig. 6A). Kinetics of synthesis and /or recruitment of Plk4 might be different between lines during the growth phase of oocytes, potentially impacting aMTOCs biogenesis.

**Fig. 6.**
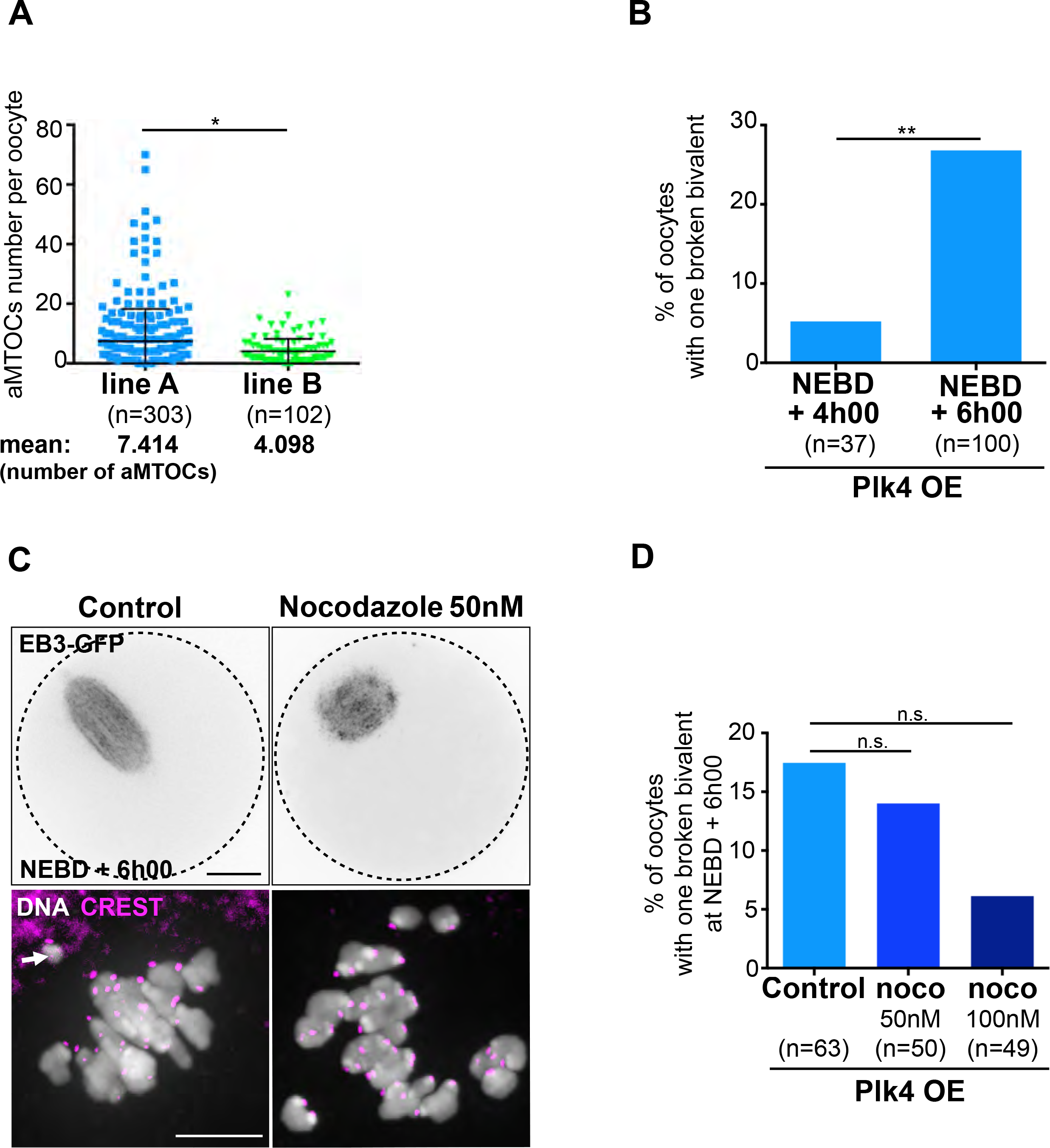
Extent of bivalent breakage correlates with aMTOCs spreading and MT density. **(A)** Quantitative analysis of aMTOCs number per oocyte in the two Plk4 OE lines (line A, blue squares, and line B, green triangles) in prophase I. p=0.0365 (two-tailed Mann-Whitney test). The mean number of aMTOCs per prophase I oocyte is indicated below the graph for each line. (**B**) Extent of bivalent breakage increases significantly as meiosis I progresses in oocytes from line A. Percentage of Plk4 OE oocytes from line A with one broken bivalent as a function of the stage of meiosis I (NEBD + 4h00 and NEBD + 6h00). p=0.0047. **(C)** Effect of the nocodazole treatment observed at NEBD + 6h00 in living Plk4 OE oocytes from line A, expressing EB3-GFP (upper panels) and in fixed samples (lower panels) labelled for DNA (white) and CREST (pink). The black dotted circles in the upper panels highlight the oocyte contour. Scale bar is 20 μm. **(D)** Decreasing MT density with nocodazole reduces the extent of DNA breakage. Percentage of Plk4 OE oocytes from line A presenting one broken bivalent at NEBD + 6h00 as a function of nocodazole concentration (0, 50 and 100nM, respectively, darker blue histograms). p=0.7966 for 0 vs 50 nM and p=0.0886 for 0 vs 100nM. For **(B)** and **(D)**, the statistical tests used are Fisher’s exact tests. For **(A**), **(B**) and (**D**), n is the number of oocytes.

We then counted oocytes presenting a broken chromosome at different time points in meiosis I (in Plk4 OE from line A). At 4h00 after NEBD, we observed only 5.4% of oocytes with a broken bivalent, compared to 27.0% at 6h00 after NEBD (Fig. 6B). At this stage, oocytes already have individualized chromosomes easier to count than at 6h00 after NEBD when they are aligned on the metaphase plate (Fig S3A and Supplementary Movie S7). The percentage of oocytes that presented a broken chromosome increased significantly as meiosis I progresses, as the spindle assembles and k-fibers are formed (Bennabi et al., 2016; Breuer et al., 2010; Dumont et al., 2007; Schuh and Ellenberg, 2007).

Since MT density is increased in Plk4 OE, maybe due to the increase in the surface of nucleation from fragmented aMTOCs (Fig. 1E and 2B), we tested whether lowering MT density would impact the extent of bivalent breakage. To achieve this, Plk4 OE oocytes from line A, were incubated with very low doses of nocodazole (ranging from 50 to 100nM). As previously shown, such doses impact on spindle MT density (Fig. 6C and S6). Nocodazole treatment had a tendency to reduce the percentage of breakage events normally observed 6 h after NEBD (14.0% with 50nM nocodazole, 6.1 % with 100nM nocodazole, compared to 17.5% in control oocytes; Fig. 6D and Supplementary Movie S8). The differences were not statistically significant (Fig. 6D) but this may be due to the small proportion of oocytes with a broken bivalent, even in non-treated Plk4 OE oocytes and to the fact that treatment with such low doses of nocodazole (20 to 10 times lower than the usual 1μM dose) may have variable effects on oocytes. Altogether our results argue for a positive correlation between aMTOCs fragmentation, spindle bipolarization and the extent of DNA breakage.

### Plk4 OE oocytes progress less efficiently throughout meiosis I

We then tested if the presence of a broken bivalent would impact on cell cycle progression of Plk4 OE oocytes. We live-imaged Plk4 OE oocytes throughout meiosis I. These oocytes were delayed by almost 2h in PBE timing compared to controls (Plk4 OE, 10h50 +/− 2h00; controls, 9h05 +/−1h36; Fig. 7A). The delay was most probably due to Spindle Assembly Checkpoint (SAC) activation, since it could be rescued using the Mps1 inhibitor, reversine, as previously described (Hached et al., 2011; Kolano et al., 2012). In the presence of reversine, Plk4 OE underwent PBE with kinetics and efficiency comparable to controls (Fig. 7A). However, the maintenance of an active SAC was not due to the presence of a broken bivalent. Indeed, by following the chromosomes using histone-GFP in live cells, we could detect the presence of a small broken fragment yet anaphase I occurred on time at 9h30 (in 7 oocytes with a fragmented chromosome, 4 matured on time; Fig. 7B). This observation is fully consistent with previous work showing that drug-induced breakage of one bivalent does not induce SAC activation (Collins et al., 2015). Since we do not observe any delay in anaphase I timing in mCherry-Plk4 cRNA injected oocytes (data not shown), which display higher Plk4 levels (Fig. 2C and D), we conclude that it is not the presence of an excess active kinase but rather global perturbation of spindle assembly that delays meiosis I progression. As a consequence of this failure to arrest, some mature eggs will be produced with structural anomalies, as observed at anaphase I (Fig. 7C).

**Fig. 7.**
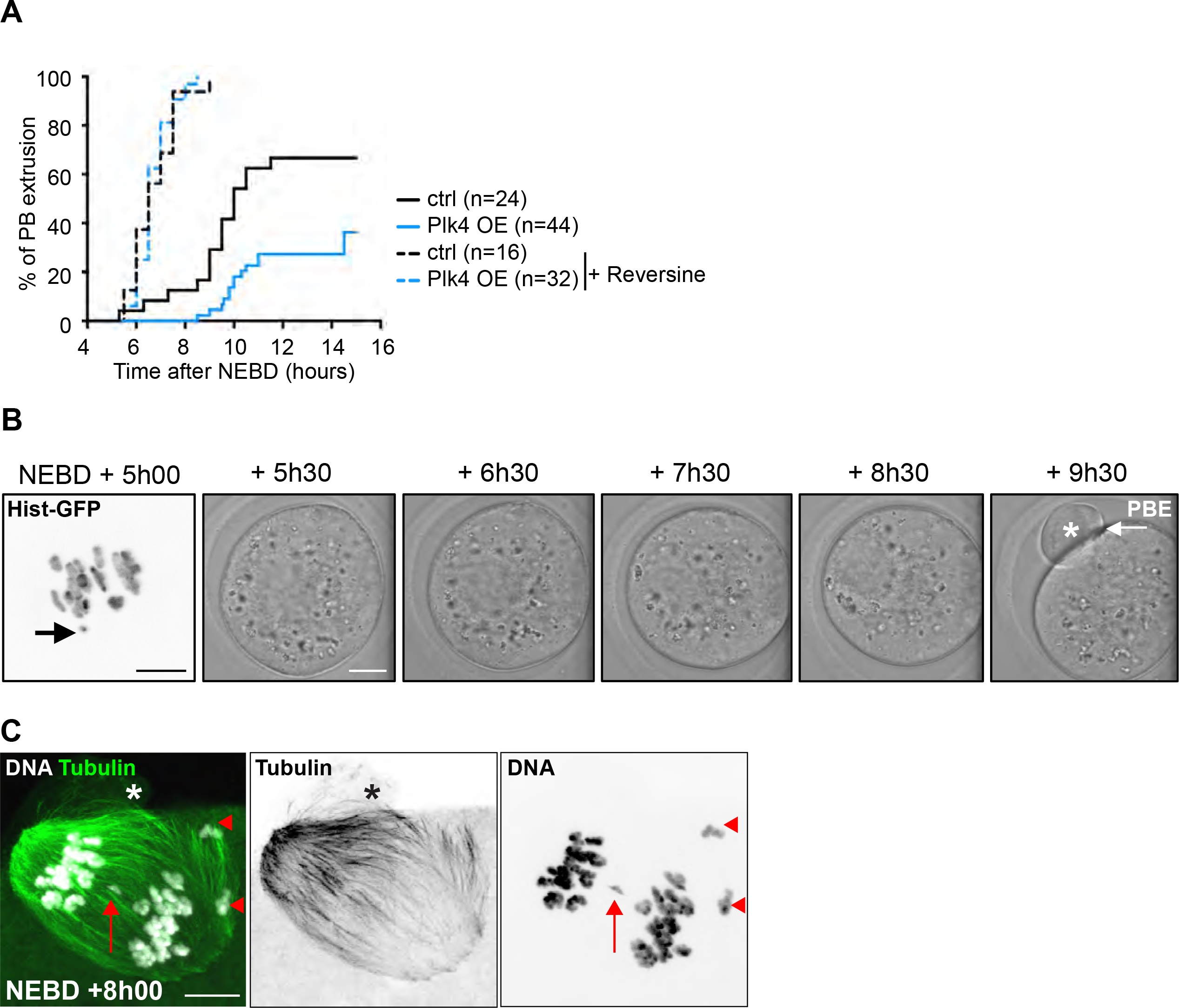
Plk4 OE oocytes have impaired progression into meiosis I due to SAC activation. **(A)** Kinetics of PBE extrusion in controls (black curves) and Plk4 OE (blue curves) oocytes from line A upon treatment (dotted curves) or not (plain curves) with 100nM Reversine at NEBD. n is the number of oocytes. PBE mean times are 9h05 for controls, 10h50 for Plk4 OE, 6h43 for controls treated by Reversine and 6h41 for Plk4 OE treated by Reversine. Tukey’s multiple comparison test gives ** for controls vs Plk4 OE, **** for controls vs controls + Reversine, controls vs Plk4 OE + Reversine, Plk4 OE vs controls + Reversine and Plk4 OE vs Plk4 OE + Reversine. Controls + Reversine and Plk4 OE + Reversine are not significantly different. **(B)** Live image of a Plk4 OE oocyte (line A) presenting a DNA fragment (chromosomes labelled with histone-GFP) which can be observed on the left picture (black arrow) and that nonetheless extrudes a first polar body at NEBD + 9h30 (PBE, Polar Body Extrusion; white arrow and white star). Scale bar is 10 μm for the left panel and 20 μm for the other panels. Time-points are in hours and minutes after NEBD. The emerging first polar body is highlighted by a white star. **(C)** Evidence for structural defects in chromosome segregation in Plk4 OE oocytes. Plk4 OE oocyte from line A, observed at NEBD + 8h00, undergoing anaphase I where DNA (white) and MT (green) are labelled. We can clearly observe the smallest DNA fragment lagging in between the two sets of univalents (red arrow) and the largest DNA fragment being separated into two deleted univalents, each retained in the oocyte (two red arrowheads). The stars indicate the position of the polar body starting to be emitted. Scale bar is 10 μm.

## Discussion

Fragmentation of aMTOCs at NEBD is normally concomitant with a burst in local MT assembly around chromosomes (Dumont et al., 2007; Schuh and Ellenberg, 2007). Here we found that precocious fragmentation of aMTOCs in conditions of Plk4 overexpression was associated with increased MT polymerization at NEBD. We showed that fragmentation of aMTOCs significantly increases their surface area, and so propose that augmentation of their capacity to polymerize MT is related to this surface area increase, providing larger sites for *γ*-TuRC recruitment. Previous work has shown that MT density controls the timing of first meiotic spindle bipolarisation (Dumont et al., 2007; Breuer et al., 2010; Kolano et al., 2012). Consistent with this, Plk4 OE oocytes displayed accelerated rates of spindle bipolarisation. In mouse oocytes, the aMTOCs morphology appears adapted to mediate the nucleation of an appropriate amount of MT at a given time and we show here that alterations from this timely regulation have deleterious consequences on meiosis outcome.

We observe that spindles assembled with higher MT density at NEBD have the capacity to induce breakage of three different chromosomes (9, 12 and 15) in different chromosomal regions; near centromeres or close to telomeres (Fig. S4C). The capacity of spindle MT to exert sufficient tension on chromosomes to result in their breakage is still a matter of debate (Ganem and Pellman, 2012). However, here we provide evidence that this can happen in meiosis in regions where a Cre-mediated recombination event may have generated a region of fragility. The mechanism at the origin of this fragility remains elusive. Yet, the break did not occur before meiosis resumption since Plk4 OE oocytes at all stages of their growth from both transgenic lines displayed overexpression of the mCherry-Plk4 transgene (data not shown). This data argues that Cre-mediated recombination and repair occurred in all oocytes, at the beginning of their growth phase, when the ZP3-Cre turns on (Lewandoski et al., 1997) to allow Plk4 transgene expression. Furthermore, the percentage of oocytes presenting one broken bivalent increased significantly (5x) from 4h00 to 6h00 after NEBD (Fig. 5B). Our data thus show that most breaks take place while the meiosis I spindle is being assembled (Fig. S7). Meiosis I spindle assembly in both mouse and human oocytes is progressive, starting with the set-up of bipolarity, then K-fiber assembly around NEBD +5h00 and ending with spindle poles assembly (Bennabi et al., 2016; Breuer et al., 2010; Dumont et al., 2007; Holubcová et al., 2015; Kitajima et al., 2011; Schuh and Ellenberg, 2007). Even though forces during the different critical steps of meiosis I spindle assembly have never been measured directly, we can speculate that these forces increase as more bundles are made and K-fibers become progressively solidly anchored into the two spindle poles. This could potentially explain the increase in breakage events observed in our model.

In mitosis, the consequences of Plk4 over-expression have been extensively studied in various model systems (Habedanck et al., 2005; Marthiens et al., 2013; Rodrigues-Martins et al., 2007), but with no reported evidence for Plk4-induced chromosome breaks. Furthermore, whole genome sequencing of Plk4 OE cells from several mouse tissues have not identified acentric chromosomes (Levine et al., 2017; Serçin et al., 2016). One main difference between previous studies and ours is that in mitotic spindles, forces are mostly exerted on kinetochores whereas in meiosis, spindle forces exerted on kinetochores are transmitted to chiasmata and to chromosome arms^13,34^. This suggests that the specific architecture of meiosis I chromosomes, bivalents held by chiasmata, is more susceptible to break under potentially increased spindle forces, in the condition of fragility during meiosis (Fig. 5E and 6F). Yet, another important difference between meiosis and mitosis, is the excessive duration of meiosis I in mouse (8 to 9 h) and human (up to 17 h) oocytes (Bennabi et al., 2016; Holubcová et al., 2015) compared to mitosis (2 h). Indeed, it is known that weak forces applied repetitively can efficiently break DNA *in vitro* (Liang et al., 2013). The extended length of meiosis I of mammalian oocytes might favour the occurrence of such breaks (Fig. S7).

Importantly, the presence of these breaks, in particular those producing acentric chromatin fragments not retained in the spindle (in Plk4 OE from line B and in MyoX), will not be detected by the SAC as the SAC functions through detecting MT occupancy at the kinetochore. Hence kinetochore-free chromatin is not recognized by any component of this checkpoint pathway. The other type of break we describe in line A will not be sufficient to elicit a robust SAC response (our results and (Collins et al., 2015)). Both types will lead to a class of chromosome abnormalities in the resulting egg that have not been observed during meiosis I before without prior treatment with DNA damaging agents. Yet such types of structural anomalies, where terminal small pieces of chromosomes are missing, are quite frequent, from 1/20000 to 1/50000 for the Wolf-Hirschhorn syndrome (Ho et al., 2016). It would be interesting to investigate the status of Plk4 activity in Wolf-Hirschhorn syndrome as affected children are precisely characterised by mutations in the Plk4 gene (Martin et al., 2014). Even though we describe here a very artificial system, our work might shed new light on the aetiology of human diseases associated to terminal deletions.

## Acknowledgements

Authors wish to thank Ludovic Dumont and Anchi Chang, two master students for their help at the beginning of the project, Carole Pennetier and Manon Chartier at the Curie Institute for genotyping, Adel Al Jord from our lab for critical reading of the manuscript, Lucie Sengmanivong from the Nikon Imaging Centre @ Institut Curie-CNRS (Paris, France), Guillaume Halet (IGDR, Rennes, France) for the gift of the HistoneH2B-GFP plasmid, Andrew Holland (Johns Hopkins School of Medicine, Baltimore, USA) for the gift of the anti-Plk4 antibody and Katia Ancelin and Edith Heard at the Curie Institute for the DNA-FISH protocol. ML was supported by a foundation ARC fellowship. This work was supported by a grant from the *Ligue Nationale Contre le Cancer* (EL/2012/LNCC/MHV), by the ANR (ANR-DIVACEN to MHV & RB, N°14-CE11), by an FRM Label (DEQ20150331758 to MHV) and by an Inca grant (PLBIO 2016-270-TRAN). This work has received support from the Fondation Bettencourt Schueller, support under the program «Investissements d’Avenir» launched by the French Government and implemented by the ANR, with the references: ANR-10-LABX-54 MEMO LIFE, ANR-11-IDEX-0001-02 PSL∗ Research University. The work was also supported by BBSRC funding to the KTJ (mechanisms of DNA damage and repair in mature oocytes, BBSRC, BB/L006006/1) and AICR now World Wide Cancer (13-0170) to VM and RB.

## Conflict of Interest

The authors declare that they have no conflict of interest.

## Methods

### Mouse strains and genotyping

*Zp3-Cre* [C57BL/6-Tg(Zp3-cre)93Knw/J] breeding pairs were obtained from Jackson Laboratories. mCherry-Plk4^flox/wt^ mice were generated by random insertion of a pCAG-loxCATlox-mCherryPlk4SV40pA construct in the genome of C57BL/6N mice (Marthiens et al., 2013). Two transgenic lines were amplified from two different founder lines with two different insertion sites and different transgene copy number and were referred, for simplicity, as line A and B. Line A corresponds to the F3 line in (Marthiens et al., 2013). Unless specified, most experiments were performed on line A, which presented a higher extent of aMTOCs fragmentation and DNA breakage. For detection of Cre-mediated multiple stop codon excision, genotyping was performed using the following primers: for the Cre 5′-GCG GTC TGG CAG TAA AAA CTA TC-3′ and 5′-GTG AAA CAG CAT TGC TGT CAC TT-3′ and for the mCherry 5′-CGC CAC CAT GGT GAG CAA GGG- 3′ and 5′-CTC GTC CAT GCC GCC GGT GG- 3. Control oocytes came either from Plk4^flox/wt^; Cre^-^ or Plk4*^wt/wt^*; Cre^+^ female mice, while Plk4 OE oocytes came from Plk4^flox/wt^; Cre^+^ female mice. In Fig. 6, we also used oocytes coming from a cross of the Plk4 line A^flox/wt^; Cre^+^ with the MyoX^flox/wt^; Cre^-^ mice. The MyoX^flox/wt^; Cre^-^ have been constructed by Genoway by insertion of two Lox P sites between exons 26 and 30 of the Myosin X endogenous locus (Fig. S4B).

### Transgene integration sites identification

TLA and next-generation sequencing, made by Cergentis, were used to determine the transgenic ssODNs integration sites as previously described (de Vree et al., 2014).

### Oocyte collection and culture

Oocytes were collected from 8- to 12-week-old female mice into M2 + BSA medium supplemented with 1 μM milrinone (Reis et al., 2006) to ensure a block in prophase I. Resumption of meiosis was triggered by culturing oocytes in milrinone-free medium. All drugs were stored in DMSO at −20°C and diluted in M2 + BSA.

### Plasmids construction and *in vitro* transcription of cRNA

We used previously described constructs: pRN3-H2B-RFP, pRN3-EB3-GFP (Breuer et al., 2010) and pMDL-Mad2-YFP (Lane et al., 2012). Histone2B-GFP (a gift from Guillaume Halet, IGDR, Rennes, France) was subcloned into the pRN3 vector. The mCherry-Plk4 construct used to produce the transgenic lines (Marthiens et al., 2013) was subcloned into the pRN3 vector. All cRNAs were synthesized using the T3 mMessage mMachine Kit (Ambion) and resuspended in RNase-free water as previously described (Verlhac et al., 2000a).

### Microinjection

Injection of *in vitro*-transcribed cRNAs into the cytoplasm of prophase I-arrested oocytes was performed using an Eppendorf Femtojet microinjector as described (Verlhac et al., 2000b) and the oocytes were further kept for 1-3 h in milrinone to allow expression of fusion proteins. Oocytes were then released from their prophase I arrest by transferring and washing into milrinone-free M2 medium.

### Drug treatments

Both nocodazole and reversine were added at NEBD. Oocytes were collected in M2 medium + Milrinone and then washed in Milrinone-free M2 medium containing either 50 nM or 100 nM of nocodazole or 100 nM reversine as in Refs (Hached et al., 2011; Kolano et al., 2012). Oocytes are kept in nocodazole until fixation at NEBD + 6h00. Control oocytes were treated with DMSO at the same dilution as the drugs.

### Live Imaging

Spinning disk movies were acquired using a Plan APO 40×/1.25 N.A. objective on a Leica DMI6000B microscope, enclosed in a thermostatic chamber set at 37 °C (Life Imaging Services), and equipped with a CoolSnap HQ2/CCD-camera (Princeton Instruments) or EMCCD camera (Evolve) coupled to a Sutter filter wheel (Roper Scientific) and a Yokogawa CSU-X1-M1 confocal scanner. MetaMorph software (Universal Imaging) was used to collect the data.

### Immunofluorescence

For MT observations, oocytes were fixed using glutaraldehyde as previously described (Terret et al., 2003). For Plk4 endogenous labelling, oocytes were fixed using ice-cold (−20°C) methanol for 10 min. For CREST staining, oocytes were fixed at 30°C for 30min in PHEM (PIPES, HEPES, EGTA, MgCl_2_) buffer containing 2% formaldehyde and 0.05% Triton-X100, and were then permeabilized for 10min in PBS containing 0.05% Triton-X100. Oocytes were extensively washed with PBS buffer between solutions. Oocytes were incubated at 4°C overnight in PBS supplemented with Tween-20 before primary antibody incubation (CREST, Abcam, 1:50). We used a rat monoclonal antibody against tyrosinated α-tubulin (YL1/2, Abcam) at 1:200 and mouse monoclonal anti-Pericentrin (BD) at 1:400. For Plk4 labelling a rabbit anti-Plk4 antibody was used at 1:500 (gift of Andrew Holland, John Hopkins School of Medicine). After primary antibodies incubation and subsequent washes, oocytes were labelled with the corresponding secondary antibodies (Jackson ImmunoResearch Laboratories, Inc.). Coverslips were mounted in Prolong Gold with DAPI (Life Technologies). SIM samples were treated with lower concentrations of antibodies to avoid signal saturation and mounted in Prolong Gold without DAPI.

Image acquisition of fixed oocytes was carried out on the SP5/AOBS confocal microscope equipped with a Plan APO 63×/1.4 N.A. objective or on the Spinning disc used for live imaging and deconvolution was applied using the Huygens software (SVI).

SIM (Structured Illumination Microscopy) acquisitions were performed on workstations of the Nikon Imaging Centre of the Curie Institute. Acquisitions were performed in 3D SIM mode with an n-SIM Nikon microscope before image reconstruction using the NIS-Elements software based on (Schermelleh et al., 2008). The system is equipped with an APO TIRF100 1.49 NA oil immersion and an EMCCD DU-897 Andor camera.

### DNA-FISH

Oocytes were cultured in M2 medium and fixed once in 1X PBS, 1% paraformaldehyde (PFA) and 1mg ml^−1^ PVP, for 1 min at RT. Cells were permeabilised at RT for 1 min in 1X PBS, 0.4% Triton X-100 and 0.5% PFA. Cells were further fixed for 10 min at RT in 1X PBS, 4% PFA, 1mg ml^−1^ PVP and 0.05% Triton X-100. Cells were then permeabilised in 1X PBS, 0.5% Triton X-100, 1mg ml^−1^ PVP and 0.02% RNAse for 1h at 37°C. Samples were pre-incubated in 1μg ml^−1^ Cot-1 DNA, 10% Dextran sulfate, 2X SSC, 0.5 mM EDTA, 0.05% Triton X-100, 1mg ml^−1^ PVP, 0.5 mg ml^−1^ BSA and 4 M urea (pH=7.3) overnight at 37°C, then denatured at 83°C for 10 minutes and pre-hybridized for 1h at 37°C. Here urea has been used instead of formamide (Sinigaglia et al., 2017). After overnight hybridization with a chromosome paint against chromosome 9 (Metasystem, see supplier’s protocol) at 37°C, oocytes were washed at 45°C, twice for 15 min in 0.2 x SSC, 0.05% Triton X-100 and 1mg ml^−1^ PVP and mounted in Prolong Gold with DAPI (Life Technologies). Denaturation and hybridization steps were performed in glass-bottom petri dishes. Other steps were performed in 4% agarose and 9 mg ml^−1^ NaCl coated petri dishes.

### Image analysis

3D analysis of aMTOCs and chromosome fragments morphology was performed using Imaris (Bitplane). Comparison of mCherry-Plk4 overexpression levels in cRNA-injected oocytes versus transgenic oocytes was performed in Metamorph by measuring the absolute intensity of fluorescence on summed Z-projections (70 Z, spaced every 0.5 μm, arbitrary units). Background values were measured within a region of interest outside the cell and were subtracted before quantification. Plk4 and Pericentrin levels in Plk4OE oocytes over endogenous protein in control oocytes were quantified on summed Z-projections (70 Z, spaced every 0.5 μm, arbitrary units). SIM acquisitions were treated in Fiji after reconstruction and realignment then rendered in Imaris 3D viewer after calibration. The total and local MT fluorescence signal intensities were measured in oocytes expressing EB3-GFP on summed Z-stack projections (6 Z planes spaced every 4 μm). Timing of bipolarization was assessed in oocytes expressing EB3-GFP imaged at 10 min intervals. Statistical analyses were performed using Graphpad Prism 5.0 software. ns, non-significant (p> 0.05); * p<0.05; ** p<0.01; *** p<0.001; **** p<0.0001.

## Supplemental Figure Legends

**Fig. S1 Quantification of the Pericentrin and Plk4 signals in the whole oocyte (A)** Measure of Pericentrin signal intensity in the whole oocyte from control (black dots) and Plk4 OE (blue squares) as observed in Fig. 1A. p=0.2277. **(B)** Measure of Plk4 fluorescence intensity in the whole oocyte from control (black dots) and Plk4 OE (blue squares) as observed in Fig. 1A. p=0.1712. Statistical tests used are two-tailed t-test with Welch correction. n is the number of oocytes.

**Fig. S2 SIM acquisition of aMTOCs in Plk4 OE oocytes suggests fragmentation at the end of oocyte growth (A)** SIM images of aMTOCs from incompetent (left) and competent (right) Plk4 OE oocytes observed in prophase I where Plk4 appears in pink and Pericentrin in green. Upper panels correspond to one Z-section acquired with SIM. Middle and lower panels correspond to surface rendering views (X/Y views for middle and Z/X views for lower panels) of the aMTOCs reconstructed in 3D using Imaris. The white arrows point toward internal cavities within the Pericentrin labelling in incompetent Plk4 OE oocytes. Scale bar is 1 μm. (**B**) Schematic representation of premature fragmentation of aMTOCs (pink) around the nucleus (in gray) in competent prophase I Plk4 OE oocytes.

**Fig. S3 Plk4OE oocytes present accelerated rates of spindle bipolarisation as observed in fixed oocytes (A)** Immunofluorescent staining of control (left panels) and Plk4 OE (right panels) oocytes fixed at NEBD + 4h00 and NEBD + 6h00. Tubulin is green, DNA is white, Pericentrin is pink. Scale bar is 10 μm. **(B)** Percentage of bipolar spindles in control (black histogram) and Plk4 OE (blue histogram) oocytes fixed at NEBD + 4h00. p= 0.0047 (Fisher exact test). n is the number of oocytes. (**C**) Immunofluorescence images from control (Plk4 line A^wt/wt^; Cre^+^ and Plk4 line A^flox/wt^; Cre^-^) and cRNA injected oocytes observed at NEBD + 6h00. DNA is in gray and CREST staining in magenta. The percentage of oocytes without any broken bivalent is indicated on the pictures. n is the number of oocytes.

**Fig. S4 Scheme illustrating the different genetic insertions used in the study (A)** Scheme of the mCherry-Plk4 transgenic construction. **(B)** Scheme of the two Lox P insertions in the endogenous Myosin X locus. **(C)** Table illustrating genomic locations of three different Lox P insertion sites as well as the percentage of oocytes presenting one broken bivalent, probably at the insertion site schematized in the middle panel, coming from the respective genotypes Plk4 line A^flox/wt^; Cre^+^, Plk4 line A^flox/wt^; Cre^+^; MyoX^flox/wt^, Plk4 line B^flox/wt^; Cre^+^. n is the number of oocytes.

**Fig. S5 Amount of mCherry-Plk4 on aMTOCs is comparable in line A and B in prophase I oocytes (A)** mCherry-Plk4 signal (pink) around the nucleus observed in live prophase I of Plk4 OE oocytes from line A (upper image) and line B (lower image). Scale bar is 10 μm. **(B)** Quantification of the mCherry-Plk4 signal observed in **(A)** at aMTOCs. p=0.986 (two-tailed Mann-Whitney). n is the number of oocytes.

**Fig. S6 Reduction of MT density in Plk4 OE oocytes after 50 and 100nM nocodazole treatment at NEBD** Quantification of the relative MT density measured at NEBD around chromosomes in Plk4 OE oocytes injected with EB3-GFP in control conditions (Control) or after treatment with 50 or 100nM nocodazole (noco). The EB3-GFP signal was measured in a Ø20 μm region around the chromosomes; total signal intensity was measured in a Ø70 μm region covering the whole oocyte. Normalized signal intensity is the ratio of local/total intensity measured for individual oocytes. p=0.0002 (Anova). Tukey’s multiple comparison test gives * for controls vs noco 50nM, ** for controls vs noco 100nM and * for noco 50nM vs noco 100nM.

**Fig. S7 Scheme recapitulating our model** Our data is consistent with precocious fragmentation of aMTOCs in Prophase I induced by Plk4 OE during oocyte growth. This in turn increases the density of MTs at NEBD, which promotes the breakage of one bivalent in any Cre-recombined Lox P-containing insertion during meiosis I. In the schemes, aMTOCs are in light pink, bivalents appear in white, microtubules in green, kinetochores in dark pink. On the pictures, DNA appears white, Pericentrin in pink and microtubules in green. The broken bivalent is highlighted in blue both on scheme and pictures.

## Supplementary Movies

**Movie S1: Spindle bipolarization is accelerated in oocytes overexpressing Plk4** Early steps of spindle formation in control (left) versus Plk4 OE oocytes (right) expressing EB3-GFP. Oocytes were followed from prophase I exit until spindle bipolarization. EB3-GFP appears white and mCherry-Plk4 magenta. Timing (expressed in hours and minutes) is relative to NEBD. Movie related to Fig. 2D.

**Movie S2: Chromosome fragment moving back and forth across the metaphase plate in Plk4OE oocyte** Chromosomes labelled with Histone-GFP followed in Plk4 OE (line A) during meiosis I. Asterisks show the small chromosome fragment. Chromosomes appear in gray levels. Timing (expressed in hours and minutes) is relative to NEBD. Movie related to Fig. 3A.

**Movie S3: 20 bivalents can be observed in control oocytes at NEBD +6h00** Z stack (Z-spacing: 200 μm) from a control oocyte observed at NEBD +6h00, stained for DNA with Dapi (white) and kinetochores with CREST (magenta). Deconvolution of the stack of images has been performed as described in the Methods section.

**Movie S4: 21 pieces of DNA can be counted in Plk4 OE line A oocytes at NEBD +6h00** Z stack (Z-spacing: 200 μm) from a Plk4 line A^flox/wt^; Cre^+^ oocyte observed at NEBD +6h00, stained for DNA with Dapi (white) and kinetochores with CREST (magenta). Deconvolution of the stack of images has been performed as described in the Methods section.

**Movie S5: 21 pieces of DNA can be counted in Plk4 OE line B oocytes at NEBD +6h00** Z stack (Z-spacing: 200 μm) from a Plk4 line B^flox/wt^; Cre^+^ oocyte observed at NEBD +6h00, stained for DNA with Dapi (white) and kinetochores with CREST (magenta). Deconvolution of the stack of images has been performed as described in the Methods section.

**Movie S6: A new form of DNA break can be observed in Plk4 line A^flox/wt^; Cre^+^; MyoX^flox/wt^ oocytes at NEBD +6h00** Z stack (Z-spacing: 200 μm) from a Plk4 line A^flox/wt^; Cre^+^; MyoX^flox/wt^ oocyte observed at NEBD +6h00, stained for DNA with Dapi (white) and kinetochores with CREST (magenta). Deconvolution of the stack of images has been performed as described in the Methods section.

**Movie S7: Bivalents can be easily counted in Plk4 OE oocytes observed at NEBD +4h00** Z stack (Z-spacing: 200 μm) from a Plk4 line A^flox/wt^; Cre^+^ oocyte observed at NEBD +4h00, stained for DNA with Dapi (white) and kinetochores with CREST (magenta). Deconvolution of the stack of images has been performed as described in the Methods section.

**Movie S8: Bivalents can be easily counted in Plk4 OE oocytes treated with 50 nM nocodazole and observed at NEBD +6h00** Z stack (Z-spacing: 200 μm) from a Plk4 line A^flox/wt^; Cre^+^ oocyte treated with 50 nM nocodazole and observed at NEBD +6h00, stained for DNA with Dapi (white) and kinetochores with CREST (magenta). Deconvolution of the stack of images has been performed as described in the Methods section.

